# Evidence for minimal cardiogenic potential of Sca-1 positive cells in the adult mouse heart

**DOI:** 10.1101/404038

**Authors:** Lauren E. Neidig, Florian Weinberger, Nathan J. Palpant, John Mignone, Amy M. Martinson, Daniel W. Sorensen, Ingrid Bender, Natsumi Nemoto, Hans Reinecke, Lil Pabon, Jeffery D. Molkentin, Charles E. Murry, Jop H. van Berlo

## Abstract

**Background:** Despite modern pharmacotherapy, heart failure remains a major medical burden. The heart has a limited regenerative capacity, and bolstering regeneration might represent new therapeutic approaches for heart failure patients. Various progenitor cells in the heart have been proposed to have cardiomyogenic properties, but this evidence is based mostly on cell culture and transplantation studies. One population of interest is characterized by the expression of Stem Cell Antigen-1 (Sca-1). Here we tested the hypothesis that Sca-1^+^ cells are endogenous progenitors for cardiomyocytes in the adult heart.

**Methods:** We evaluated the innate cardiogenic potential of Sca-1^+^ cells *in vivo* by generating a novel mouse model to genetically lineage-trace the fate of Sca-1 expressing cells. This was accomplished by introducing a tamoxifen-inducible Cre-recombinase into the Sca-1 locus (Sca-1^mCm/+^). Crossing this mouse line to a Cre-dependent tdTomato reporter line allowed for genetic lineage-tracing of endogenous Sca-1^+^ cells (Sca-1^mCm^R26^tdTomato^). The frequency of Sca-1^+^ cardiomyocytes was quantified from dispersed cell preparations and confirmed by in situ histology.

**Results:** We validated the genetic lineage tracing mouse model in bone marrow and heart. Unlike previous publications suggesting significant cardiogenic potential, we found that less than 0.02% of cardiomyocytes per year were derived from Sca-1^+^ cells in the adult heart under homeostatic conditions. At six months after myocardial infarction, we found less than 0.01% of cardiomyocytes were derived from Sca-1^+^ cells.

**Conclusion:** Our results show that Sca-1^+^ cells in the adult heart have minimal cardiogenic potential under homeostatic conditions or in response to myocardial infarction.

## INTRODUCTION

Cardiac regeneration has been investigated for more than 100 years, and the ability of the adult heart to generate new myocytes has been under debate ever since(1, 2). The current consensus is that *de-novo* cardiomyogenesis is limited to ∼1% per year in the healthy adult mammalian heart(3, 4). The extent by which this limited ability to generate cardiomyocytes is increased after injury remains to be determined. However, even 1% per year *de novo* cardiomyogenesis might represent a target for regenerative therapeutic interventions. Whether newly-formed cardiomyocytes are derived from cellular division of pre-existing myocytes or from differentiation of progeny of a resident cardiac progenitor cell (CPC) population is topic of debate(5–6). Most recent evidence points towards proliferation of pre-existing myocytes as the main source of newly formed cardiomyocytes(7–9). On the other hand, there also is an extensive literature on multiple types of resident CPCs, including distinct populations that express c-Kit or Stem Cell Antigen-1 (Sca-1, reviewed in(10)). CPCs have been posited as a source of endogenous cardiomyocyte renewal and as cells that can be harvested, expanded *in vitro* and delivered therapeutically after infarction. The cardiogenic potential of these distinct CPC populations has usually been demonstrated in transplantation studies(11–13). Yet, it is increasingly recognized that transplantation studies do not always reveal the endogenous functions of stem cells(14), and more recent fate-mapping studies revealed that one putative CPC population, characterized by the expression of c-kit does not significantly contribute to cardiomyocytes in vivo(15–17), further stimulating the debate of whether a true CPC population exist.

This study tested the hypothesis that cells expressing Sca-1 are endogenous CPCs in the adult mouse heart. Sca-1 was initially described as a surface marker of murine hematopoietic stem cells(18), however, its expression is not restricted to stem cells, as it is also expressed on mature B- and T-lymphocytes as well as many cells in the thymus, brain, kidney, lung and the heart(19–21). Oh et al. were the first to describe a perivascular Sca-1^+^ cell that does not express CD45, CD34 or CD31(22), suggesting it is neither a leukocyte nor an endothelial cell. These Sca-1 positive cells, which also express PDGFRα(23), were reported to differentiate into endothelial cells, smooth muscle cells and cardiomyocytes after transplantation(22). The coexpression of Sca-1 and PDGFRα was also reported to define a mesenchymal stem cell-like population in the murine heart that demonstrated some cardiogenic potential in vitro(24). However cells sorted for PDGFRα from fetal human hearts only differentiated into smooth muscle and endothelial cells, but not cardiomyocytes(25).

To assess the ability of endogenous Sca-1^+^ cells to generate cardiomyocytes, a previous attempt was made to genetically lineage-trace Sca-1^+^ cells in the adult mouse heart(26). This study reported a frequent contribution of Sca-1 expressing cells to cardiomyocytes, although precise quantifications were not provided. A limitation of this study is their use of a short region of the Sca-1 promoter to drive expression of Cre recombinase via transgenesis. The transgene showed more widespread expression than just Sca-1 expressing cells, complicating interpretation of the results(27). Adequate specificity and sensitivity are major concerns for all fate-mapping studies and might explain contradictory results(15, 28–30). To reliably trace the descendants of Sca-1^+^ cells *in vivo*, we generated a tamoxifen-inducible Sca-1 mER-Cre-mER knock-in mouse model using a strategy that has been successfully applied in the hematopoietic system(31), and tested the hypothesis that endogenous Sca-1^+^ cells are progenitors for cardiomyocytes *in vivo* under physiological and pathophysiological conditions. Upon validating the model in hematopoietic stem cells, we report that Sca-1^+^ cells do not form new cardiomyocytes in healthy myocardium or after myocardial infarction.

## METHODS

Data, analytic methods and study material will be made available to other researchers for purpose of reproducing the results by contacting the corresponding authors.

### Animals

All animal procedures were conducted in accordance with the US NIH Public Health Service Policy on Humane Care and Use of Laboratory Animals, and all procedures complied with and were approved by either the University of Washington or the University of Minnesota Institutional Animal Care and Use Committee. Both male and female mice between 8-12 weeks of age, weighing an average of 25g, were randomly assigned to each experimental group. Mice were group housed by sex throughout the experiments in ventilated cages and were of normal immune status.

### Generation of the Sca-1 mER-Cre-meER and Sca-1^mCm^R26^tdTomato^ mouse models

We generated the knock-in mouse model adapting a strategy from Hanson et al.(31) The second exon of the Ly6a gene was targeted in W4 mouse embryonic stem cells (derived from mouse line 129S6/SvEvTac). Correct integration was confirmed with Southern blot and PCR (Suppl. Fig. 1A, B). A correctly targeted clone was aggregated with morula stage embryos, transferred into pseudopregnant mice, and chimeric offspring were back-bred onto C57Bl/6 mice. To create inducible tdTomato labeling of Sca-1^+^ cells, we cross-bred Sca-1^mCm^ mice with Cre-dependent reporter mice. *Gt(ROSA)26Sor*^*tm*^*4*^*(ACTB-tdTomato,-EGFP)Luo*^/J, (R26^mTmG^; strain 007676 The Jackson Laboratory) and *Gt(ROSA)26Sor*^*tm*^*1*^*(CAG-lacZ,-EGFP)Glh*^/J, previously cross-bred with Rosa26-Flpe mice and back-crossed to C57Bl/6 (R26^GFP^; strain 012429 The Jackson Laboratory) were initially used. Although robust recombination was seen in the heart, we observed minimal recombination in the bone marrow with this reporter strain, so we switched to (*Gt(ROSA)26Sor*^*tm*^*14*^*(CAG-tdTomato)Hze*^/J (tdTomato; strain 007914 The Jackson Laboratory) to generate most of the Sca-1^mCm^R26^tdtomato^ mice used in this study. As will be demonstrated, this reporter strain gave recombination in both the heart and hematopoietic system.

**Figure 1:**
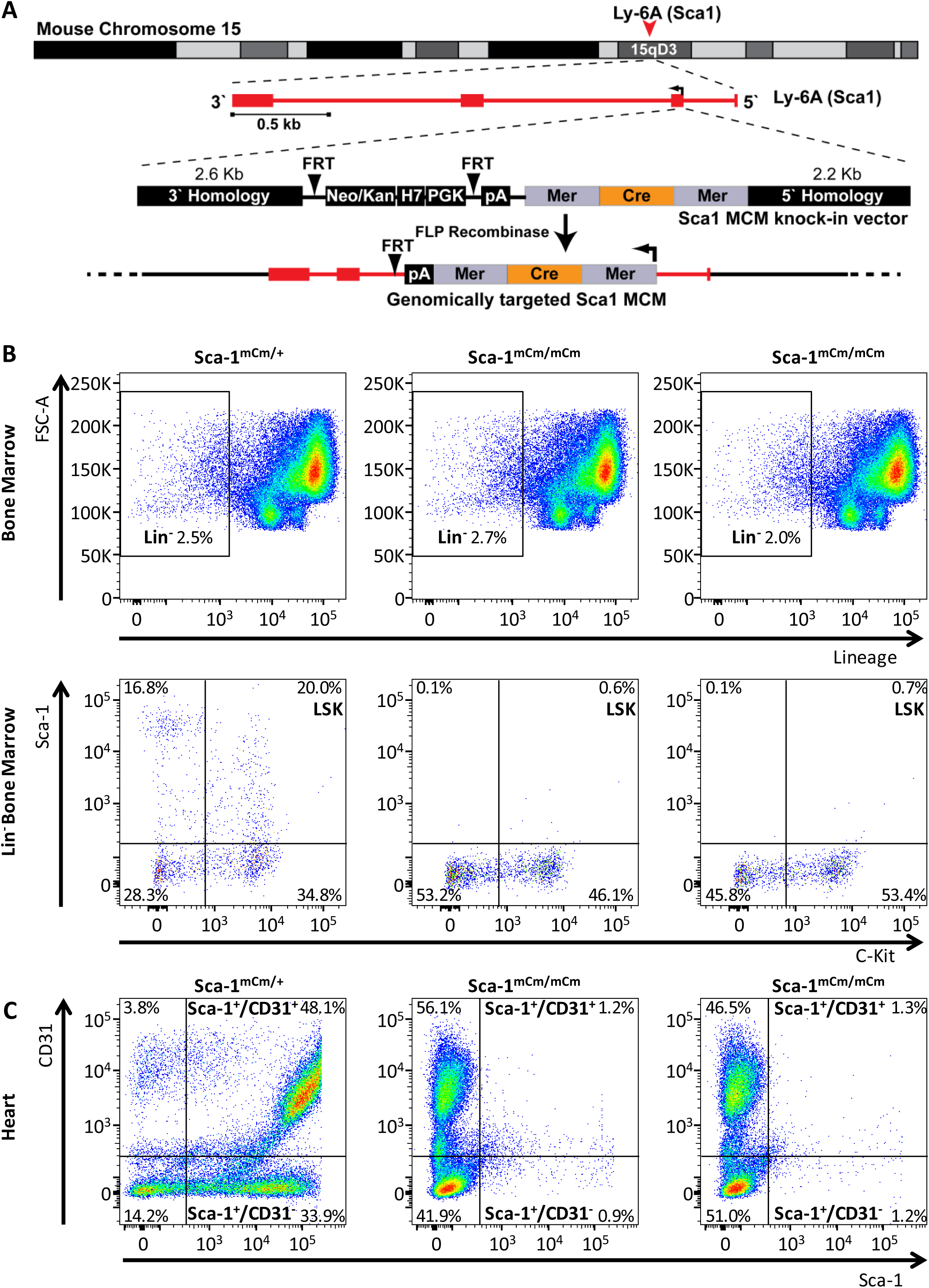
Generation of a tamoxifen-inducible Sca-1 mER-Cre-mER mouse line. A) Exon 2 of the Ly6a locus was targeted to generate the Sca-1^mCm/+^ mouse embryonic stem cell line and knock-in mice. **B**) Heterozygous Sca-1^mCm/+^ animals showed regular Sca-1 staining in the bone marrow by flow cytometry, whereas homozygous Sca-1^mCm/mCm^ did not express Sca-1 protein. This confirms correct targeting of the target locus. **C**) Heterozygous animals showed regular Sca-1 staining in the heart by flow cytometry, whereas homozygous Sca-1^mCm/mCm^ hearts did not express Sca-1 protein. Percentages indicate fraction of cells within each quadrant.

### Genotyping

1mm punch biopsies were obtained from ear pinna samples of mice and DNA was prepared using the REDExtract-N-Amp^™^ Tissue PCR Kit Protocol (Sigma-Aldrich, XNAT). PCR was performed to detect the presence of Cre recombinase using primers 5’ –ACGACCAAGTGACAGCAATG (forward) and ACCAGAGACGGAAATCCATCGCT (reverse). PCR conditions were 94 °C for 5 minutes to separate strands, followed by 30 cycles of amplification (94 °C for 30 s, 57 °C for 30 sec, 72 °C for 30 s). Additionally, knock-in specific PCR primers were used 5’-CTGCATTCTAGTTGTGGTTTGTCC (forward) and 5’-AAAGTCAAAGTCCAGGCAATAAGG (reverse).

### Animal procedures

We compared a variety of regimens to induce maximal recombination of the Cre reporter allele in the heart and bone marrow. These included the use of standard tamoxifen (oral or intraperitoneal), the biochemically active metabolite 4-hydroxy-tamoxifen, the tamoxifen analog raloxifene, and the co-administration of ketoconazole with tamoxifen to inhibit the ABC family of drug efflux pumps. We found that a 7-day course of tamoxifen yielded optimal recombination. Tamoxifen was suspended in corn oil and administered by daily intraperitoneal injections (2 mg) for 7 day to activate the inducible Cre-recombinase. Myocardial infarction surgery was performed after a seven days wash-out period. For myocardial infarction surgery, 8-12 week-old mice were anesthetized by intraperitoneal injection of ketamine and xylazine. Mice were intubated, and mechanically ventilated. Lateral thoracotomy was performed and the left anterior descending coronary artery was permanently ligated as previous described(32). Mice received buprenorphine analgesia and were monitored regularly until endpoint analysis.

### Isolation of marrow cells and non-myocytes from the heart

Bone marrow was harvested by flushing femurs and tibiae with cold Hanks Balanced Salt Solution (HBSS). Cells were centrifuged at 400 g for 10 minutes at 4°C and pellets were resuspended in HBSS. Red blood cell lysis was performed with a lysis buffer containing NH_4_Cl (155 mM), NaHCO_3_ (12 mM) and EDTA (0.1 mM). Non-myocytes were isolated from minced adult hearts by enzymatic digestion with collagenase type II (Worthington 240U/mg) for 30 minutes at 37 °C. Cells were then filtered through a 70 µm mesh. Bone marrow and non-myocytes were stained with antibodies against Sca-1 (Milteny Biotec; clone: D7), CD31 (BioLegend, clone MEC13.3), NG2 (Millipore Sigma, AB5320) a Lineage marker cocktail (BioLegend, clone, anti-mouse CD3e, clone 145-2C11; anti-mouse Ly-6G/Ly-6C, clone RB6-8C5; anti-mouse CD11b, clone M1/70; anti-mouse CD45R/B220, clone RA3-6B2; anti-mouse TER-119/Erythroid cells, clone Ter-119), Cardiac troponin T (Thermo Scientific, clone 13-11), Myosin light chain 2 (Proteintech, 10906-1-AP) and tdTomato (Abcam, ab62341). Flow cytometry was performed using a BD FACSCanto^™^ and a FACSAria II flow cytometry system. Cells were gated by forward scatter versus side scatter to eliminate debris. A minimum of 50,000 events was counted for each analysis. Analysis of flow cytometry data was performed using FlowJo software.

### Isolation of adult mouse cardiomyocytes

Mouse cardiomyocytes were isolated using an enzymatic digestion and mechanical dispersion method previously described(33). In brief, after retrograde perfusion with calcium-free Tyrode buffer (126mM NaCl, 5.4mM KCl, 0.33mM NaH_2_PO_4_, 1mM MgCl_2_, 10mM HEPES, 10mM Glu, 20mM Taurine, 25uM Blebbistatin) and digestion with collagenase solution (collagenase type II, Worthington 240U/mg), the ventricular myocytes were separated using a fine scalpel and scissors. Afterwards cells were fixed in 4% paraformaldehyde and processed for immunohistochemical analysis.

### Immunohistochemical analysis

Histological stains and subsequent analysis were conducted as described previously by our group(34–36). In brief, hearts were perfused with PBS and 150 mM KCl solution after harvesting, fixed in 4% paraformaldehyde and dehydrated overnight in 30% sucrose solution. Sections of the heart were sliced along the transverse axis into 2-mm-thick sections, frozen in OCT, cryosectioned at 5 microns, and stained with appropriate primary and secondary antibodies. Immunofluorescent images were collected by a Zeiss Axio Imager M1 upright microscope or a Nikon A1 Confocal System attached to a Nikon Ti-E inverted microscope platform. For figure preparation, images were processed to convert colors by Nikon NIS Elements software.

### Statistical analysis

Only animals that did not survive the MI surgery (<20% mortality) were excluded from the study. Animal numbers used are reported for each experiment as well as the exact total number of tdTomato^+^ cardiomyocytes identified. Screening for tdTomato positive cardiomyocytes after MI was performed blinded by one analyzer who was unaware of the genotype/Tamoxifen status of the mouse at the screening time. A minimum of 40,000 cardiomyocytes was screened from each mouse. All statistical test were performed using GraphPad Prism 4.

## RESULTS

### Generation of Sca-1 lineage-tracing mouse and selection of reporter lines

To track the lineage of Sca-1^+^ cells in the heart, we performed traditional gene-targeting in W4 mouse embryonic stem cells and inserted a cDNA cassette encoding for tamoxifen-inducible Cre recombinase (mER-Cre-mER) into the endogenous murine Ly6a locus, which encodes for the Sca-1 protein (Fig. 1A). Proper heterozygous integration was determined by Southern blotting and PCR followed by DNA sequencing (Suppl. Fig. 1A). A correctly targeted clone was aggregated with morula stage embryos to generate chimeric mice. Chimeric mice were back-crossed onto C57Bl/6 mice to generate the Cre-driver strain, which we refer to as Sca-1^mCm/+^ hereafter. The Ly6A gene resides in the Ly6 cluster on chromosome 15, which contains multiple gene duplications resulting in related proteins and pseudogenes. This complexity has led to debate and confusion regarding, for example, expression of the allelic genes Ly6A and Ly6E *(18, 37, 38)*. To conclusively verify gene targeting of the correct Ly6 gene in the Sca-1^mCm/+^, we generated homozygous Sca-1^mCm/mCm^ mice, which showed no Sca-1 protein expression by flow cytometry on either bone marrow cells or non-myocytes from the heart (Fig.1B and 1C).

Since it is known that different responder lines have different sensitivity for Cre recombinase(39), we next screened different Cre-reporter mice to identify an optimal system for Sca-1 lineage analysis. Initially, we cross-bred Sca-1^mCm/+^ mice with Rosa26^mTmG^ mice and Rosa26^GFP^ mice. When adult mice were pulsed daily with tamoxifen for one week and studied 7 days later, there was extensive recombination in the heart (Suppl. Fig. 2A and C). Flow cytometry demonstrated that on average 35% of the cells expressing Sca-1 protein expressed GFP (data not shown). However, there was no significant recombination in bone marrow hematopoietic compartment with either reporter line, where the non-endothelial Sca-1^+^ cells were essentially negative for GFP (Suppl. Fig 2B). These results could be due to insufficient Cre fusion protein, or insufficient tamoxifen to nuclearize the Cre protein, or inefficiency of Cre mediated recombination of the reporter allele. Although increasing Cre abundance did appear to increase cardiac cell labeling (Suppl. Fig. 2E), neither the use of homozygous Sca-1^mCm/mCm^ mice, nor the use of various tamoxifen/raloxifene dosing regimens labeled hematopoietic cells in the marrow. To assess if we selected for a specific subpopulation of Sca-1^+^ cells that underwent recombination due to different promoter activities, we determined Cre recombinase expression levels in Sca-1^+^ recombined and un-recombined cells. These showed no difference in expression of the mERCremER transgene, indicating that the relative inefficiency of recombination was not due to different subsets of promoter activity in subpopulations of Sca-1^+^ cells (Suppl. Fig. 2F). We therefore tested a third reporter strain. In studies of gastric stem cells(40) the Rosa26-tdTomato mouse line was identified as a highly sensitive Cre reporter strain. Thus, we cross-bred this reporter line with Sca-1^mCm/+^ mice and now observed recombination rates in the bone marrow compartment of 0.25-2% in response to tamoxifen treatment (Fig. 2A and B). Additionally, we were able to detect a population of tdTomato^+^ cells in the bone marrow that was also Lin-positive (i.e. expressed markers for the lymphoid, erythroid or granulocytic lineages; Fig. 2C), indicating that we had induced recombination in hematopoietic stem cells, which then differentiated into mature blood cells. Importantly, there was recombination in the heart averaging 33% in Sca-1^+^ /CD31^+^ populations and 13% in the Sca-1^+^ /CD31^−^ population (Fig. 2D and 2E). Subsequent studies were therefore performed in mice obtained by crossing Sca-1^mCm/+^ driver line to the tdTomato mouse, which we refer to as Sca-1^mCm^R26^tdTomato^ mice (Fig. 2A).

**Figure 2:**
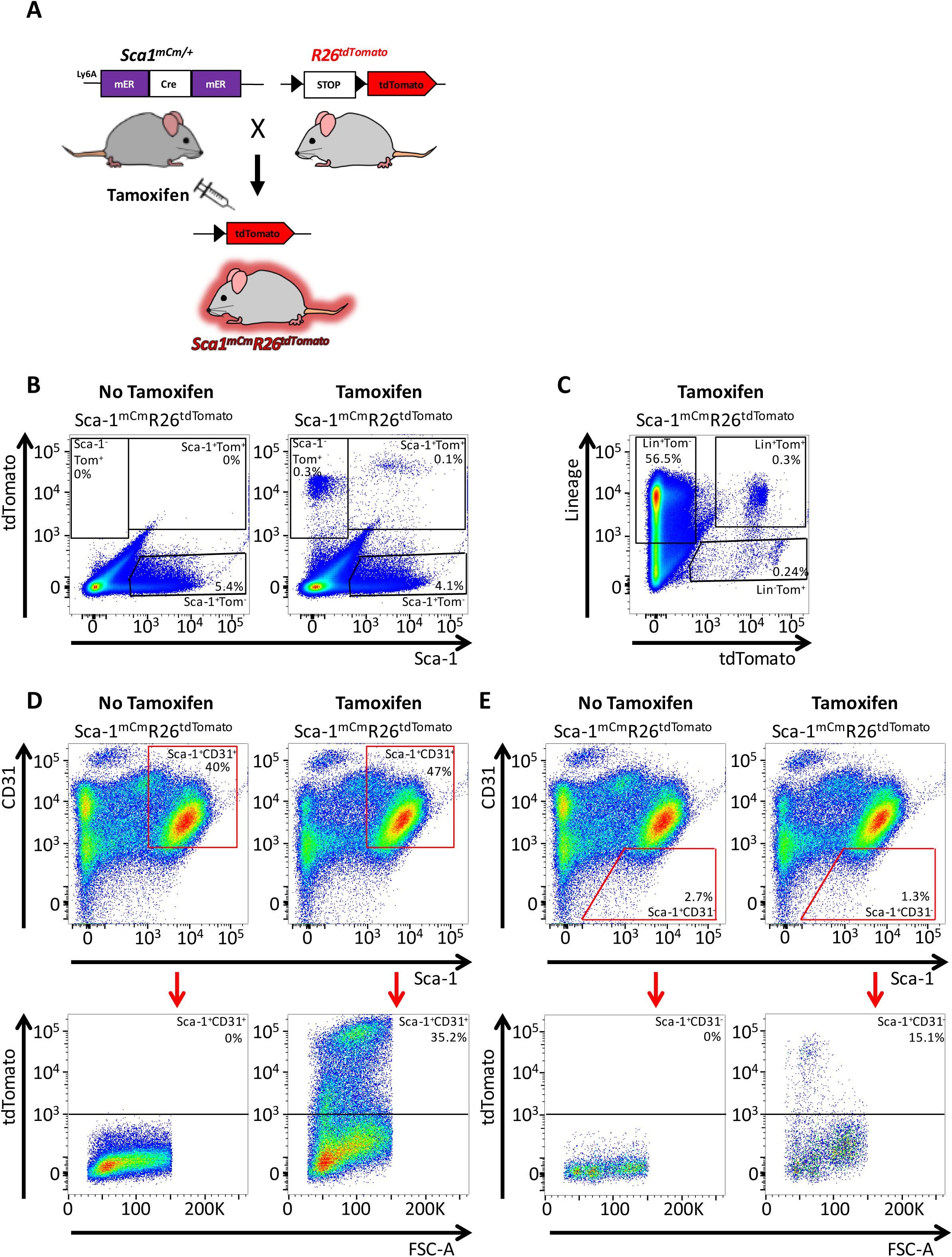
Characterization of Sca-1^mCm^R26^tdTomato^ mice. A) Heterozygous Sca-1^mCm/+^ mice were cross-bred to a Cre-dependent tdTomato reporter line and treated with tamoxifen via daily IP injections for one week. Experiments shown in B-E were performed after a one week chase period. **B)** Tamoxifen administration resulted in labeling of bone marrow (BM) stem cells (Sca-1^+^ Tom^+^) that over the pulse period differentiated into Sca-1 negative hematopoetic cells (Sca-1^−^ Tom^+^). **C)** Lineage staining revealed a population of Lin^+^ tdTomato^+^ cells (differentiated bone marrow lineages), as well as Lin^−^tdTomato^+^ cells **D)** Sca-1^+^ CD31^+^ cells isolated from Sca-1^mCm^R26^tdTomato^ hearts demonstrated successful recombination after tamoxifen induction. **E)** Sca-1^+^ CD31^−^ cells isolated from Sca-1^mCm^R26^tdTomato^ hearts demonstrated successful recombination after tamoxifen induction.

### Sca-1^+^ lineage marks microvascular endothelium and perivascular cells in the normal heart

Adult Sca-1^mCm^R26^dTomato^ animals were treated with tamoxifen daily for 7 days and hearts were harvested 1 week after the last tamoxifen injection to assess recombination by flow cytometry and histology. No tdTomato^+^ cells were identified by flow cytometry (data not shown) or by histology (Suppl. >Fig. 3) in Sca-1^mCm^R26^dTomato^ mice that did not receive tamoxifen. In mice that received tamoxifen there was a close overlap between Sca-1 protein staining and expression of tdTomato, in a pattern that suggested a vascular distribution (Fig. 3A). Co-staining for endothelial cells with CD31 antibodies and for tdTomato expression in Sca-1^mCm^R26^dTomato^ hearts demonstrated that most of the tdTomato^+^ cells were vascular endothelium (Fig. 3B). In addition, there were CD31 negative cells in the perivascular compartment that expressed abundant tdTomato. To test if these cells were pericytes, we co-stained sections for the pericyte marker NG2 and tdTomato. There was occasional co-labeling of NG2 and tdTomato, but in most instances, these were adjacent cells (Fig. 3C). TdTomato was not identified in any cardiomyocytes in tissue sections (Fig. 3D). Additionally, tamoxifen-dependent tdTomato expression in tissues known to express Sca-1 (liver, kidney, ileum, lung) confirmed the presence of tdTomato^+^ cells, predominately in an endothelial pattern, whereas tdTomato expression was absent in tissue from animals that did not receive tamoxifen (Suppl. Fig. 3).

**Figure 3:**
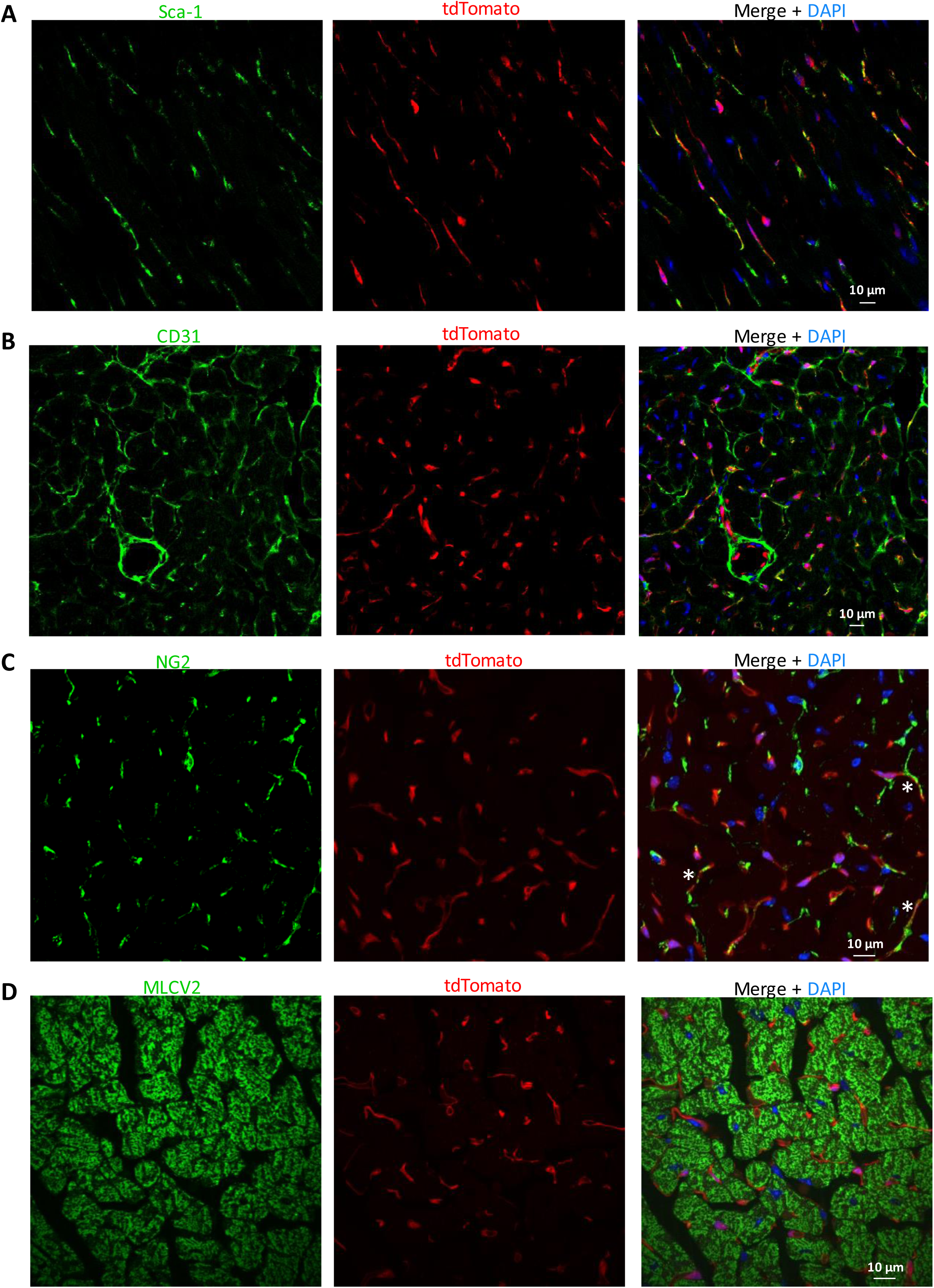
Sca-1^+^ lineage marks microvascular endothelium and perivascular cells in the normal heart. A) TdTomato expression demonstrated a vascular staining pattern that overlapped Sca-1 staining. **B**) TdTomato expression overlapped with CD31 staining. **C**) tdTomato expression sometimes overlapped with the perivascular marker NG2 as illustrated by the asterisks. **D**) TdTomato expression did not overlap with the cardiomyocyte marker MLC2V.

### Lineage tracing in normal and injured hearts reveals low numbers of Sca-1 derived cardiomyocytes

Next, we set out to test whether new cardiomyocytes were derived from descendants of Sca-1^+^ cells after myocardial infarction. Adult Sca-1^mCm^R26^dTomato^ mice were treated for 7 days with tamoxifen, while control mice were not exposed, followed by a 7-day chase period. The mice then underwent myocardial infarction, induced by permanent ligation of the left coronary artery. Mice were euthanized approximately 6 months after infarction for examination of the heart (Fig. 4A). To unambiguously determine the number of tdTomato^+^ cardiomyocytes, we enzymatically dissociated cardiomyocytes and quantified the fraction of tdTomato^+^ cardiomyocytes from live cells (uninjured heart), or after fixing and immunostaining for cardiac troponin T (cTnT) and tdTomato (post-MI). This allowed unambiguous distinction of cardiomyocytes and non-myocytes (Fig. 4D and Suppl. Fig. 4A). We first assessed the numbers of tdTomato^+^ cardiomyocytes at baseline before injury. We detected 6 tdTomato^+^ cardiomyocytes from 5 mice, averaging 0.00023% of total cardiomyocytes (Fig. 4B). Next, we assessed labeling of cardiomyocytes after myocardial infarction. Confocal line scanning in these fixed and double-stained cells demonstrated that there was no bleed-through between the troponin and tdTomato channels (Suppl. Fig. 4B). After MI, we detected only 19 tdTomato^+^ cardiomyocytes from 7 hearts. This accounts for an average of 0.007% tdTomato^+^ cardiomyocytes (Fig. 4C and Suppl. Fig. 5). Importantly, not a single cardiomyocyte expressed tdTomato in animals that were not exposed to tamoxifen, demonstrating that this is not a leaky tracing system.

**Figure 4:**
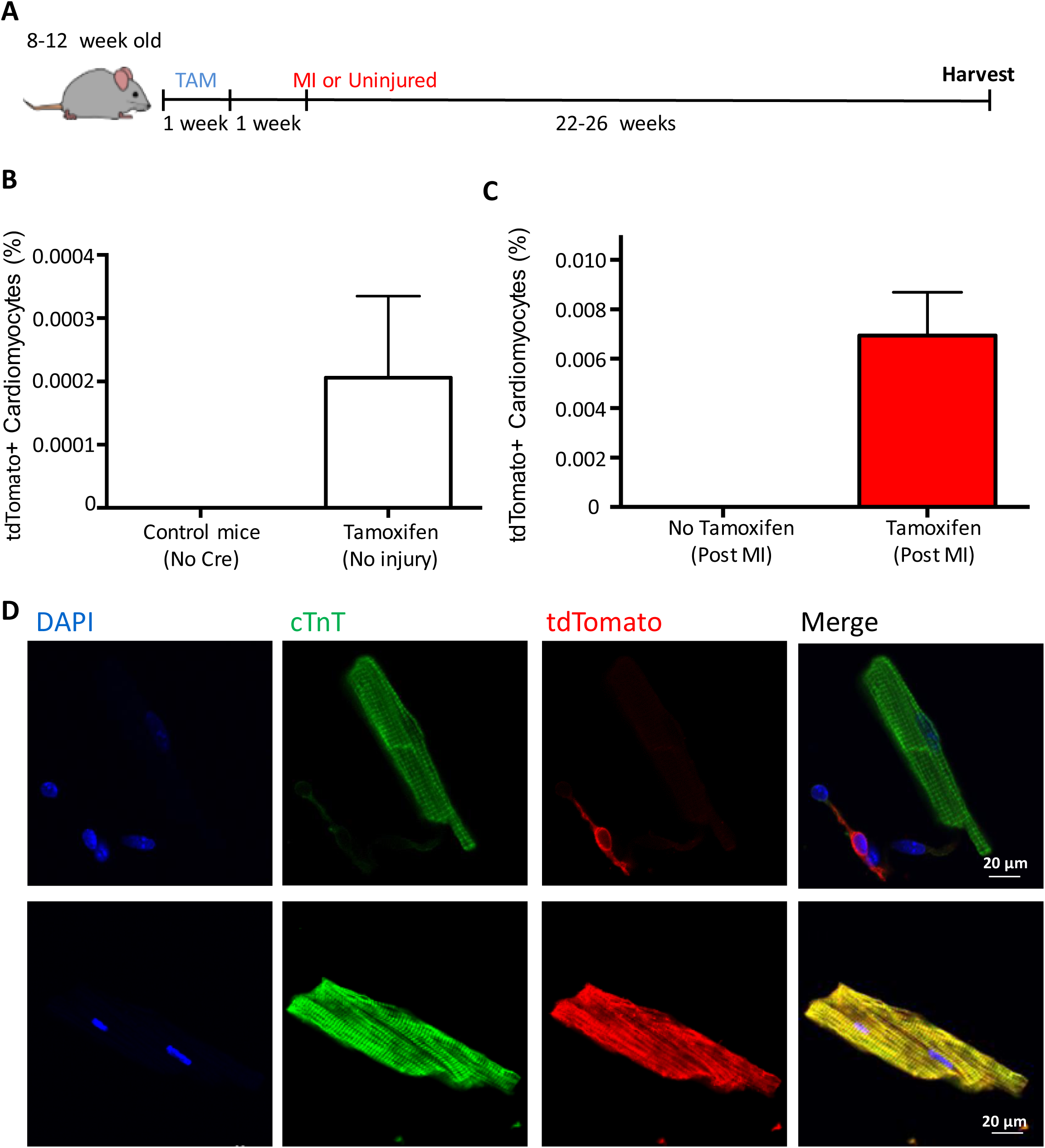
Lineage tracing in injured hearts reveals low numbers of Sca-1 derived cardiomyocytes A) Sca-1^mCm^R26^tdTomato^ animals were pulsed with tamoxifen for 1 week. LAD-ligation was performed after a wash-out period of another 7 days and hearts were harvested after 6 months. **B)** Quantification of tdTomato+ cardiomyocytes 7 days after tamoxifen from live cardiomyocytes. **C)** After MI, a total of 19 tdTomato^+^ cardiomyocytes were detected in 7 animals than underwent MI, accounting for a derived average percentage of 0.007% Sca-1^+^ - derived cardiomyocytes (Error bar = SEM). No tdTomato^+^ cardiomyocytes were observed in the absence of tamoxifen treatment. **D)** Enzymatic digestion allowed for an unambiguous differentiation between cardiomyocytes and non-myocytes. Asterisk indicates a tdTomato^+^ non-myocyte, while the cTnT^+^ cardiomyocyte in the same image is tdTomato-negative. Bottom panel: example of a rare tdTomato^+^ cardiomyocyte.

## DISCUSSION

The results of this study do not support the hypothesis that Sca-1 expressing cells have the ability to significantly contribute new cardiomyocytes to the heart after myocardial infarction in the mouse. Despite using the best available techniques for lineage tracing, including knocking an inducible Cre recombinase into the Sca-1 gene, using a sensitive Cre-reporter strain, and enzymatically dispersing cardiomyocytes from the non-myocytes to allow for single cell screening, we were unable to detect significant contributions of the Sca-1 lineage to cardiomyocytes. We conclude that Sca-1^+^ cells are not endogenous cardiac progenitor cells in the adult heart.

The results of our study contrast data from Uchida et al., who showed a significant contribution of Sca-1^+^ cells to cardiomyocytes(26). The main difference between these two studies concerns the way lineage-tracing was initiated, i.e. the Cre-driver lines. We generated a knock-in mouse line where a tamoxifen-inducible Cre-recombinase was inserted into the endogenous Ly6a locus. Uchida et al. used a transgenic approach with a 14 kb fragment of the Sca-1 gene. Transgenic approaches often demonstrate a lower fidelity of gene expression (41), while knock-in strategies often result in higher fidelity but bear the (small) risk of haploinsufficiency. The transgenic system used by Uchida et al. has been reported to generate a large number of false-positives, meaning the promoter fragment is active in cells that normally do not express Sca-1(27). Therefore, we believe the transgenic promoter fragment may have overestimated the true cardiogenic potential of endogenous Sca-1 expressing cells in this model. The Sca-1 expression pattern and the number of Sca-1^+^ cells in our Sca-1^mCm/+^ model did not differ from wild type animals arguing against haploinsufficiency. This is further corroborated by the fact that Sca-1^mCm/+^ animals did not present a cardiac phenotype (e.g. regular left ventricular function by echocardiogram, data not shown), although homozygous deletion of Sca-1 does result in mild cardiac dysfunction(42).

Our study is in line with recent work on c-kit lineage-tracing in the heart(15). C-kit^+^ cells were reported to contribute to cardiomyocytes in the adult heart(28), yet, eventually a knock-in approach that allowed to lineage trace the fate of c-kit^+^ cells *in vivo*, demonstrated that they do not contribute to cardiomyocytes in relevant numbers(15, 43). Similar to the differences between Uchida et al. and our study, the first study used a short promoter fragment that was known to have more widespread expression, and delivered this transgene via lentivirus, which likely transduced both cardiomyocytes and non-myocytes in the heart. This approach likely overestimated the true cardiogenic potential of c-kit expressing cells(28).

A different problem is the possibility of activation of Sca-1 within cardiomyocytes. Recently, it was shown that the low level of cardiomyocyte labeling in c-kit genetic lineage tracing experiments is likely not derived from differentiation of c-kit expressing cells, but rather the result of a fusion event, or due to expression of c-kit within cardiomyocytes(30). Inducible lineage tracing studies of cardiac progenitors have this inherent risk of producing *erroneous positives* due to transient Cre recombinase activity in pre-existing cardiomyocytes. Although we cannot rule out this possibility, it is likely of no relevance given the extremely low overall rate of cardiomyocyte labeling.

A third problem that might limit interpretation of genetic lineage tracing strategies is incomplete activity of Cre recombinase giving rise to *false negative* cells. We acknowledge that our recombination rates in the bone marrow were lower than expected, considering that the majority of murine hematopoietic stem cells in the bone marrow express Sca-1 protein. We evaluated several tamoxifen regimens to increase recombination rates, including the long term application of tamoxifen citrate via chow (400 mg/kg chow) for six months(15), the use of high doses of the active tamoxifen metabolite OH-tamoxifen (0.5 mg i.p. for 14 days)(44), high doses of tamoxifen (up to 320mg/kg i.p. for 5 days) or the tamoxifen alternative raloxifen (160mg/kg BW/day p.o for four weeks)(45). These strategies did not increase recombination rates. However, Cre expression did not differ between recombined vs. non recombined Sca-1^+^ cells which indicates that these did not represent two distinct cell populations in terms of Sca-1 promoter activity. The recombination rates in the heart were reasonable, but still only resulted in 0.007% of cardiomyocytes to become labeled. Thus, although this number might be a slight underestimation, it is two orders of magnitude too low to explain a 1% per year cardiomyocyte turnover rate. It is important to remember that all biological assays have some background noise, and we probably are approaching the sensitivity threshold of even the most rigorous assay when detecting only 0.007% positive cells. However, our assay was able to readily detect differentiated non-cardiomyocytes in the heart, further validating our assay. Moreover, using our rigorous approach of isolating adult cardiomyocytes from the heart to unambiguously assess tdTomato status, we did not detect a single tdTomato^+^ cardiomyocyte in animals that were not subjected to tamoxifen, indicating a lack of leakiness of the inducible Cre recombinase.

In conclusion, we did not find evidence that in the adult heart, cardiomyocytes are derived in significant numbers from Sca-1 progenitor cells. Our study echoes recent work on c-kit^+^ cells in the heart. Although there is no orthologue for Sca-1 in the human genome, the perivascular Sca-1^+^ cells of the mouse appear similar to perivascular PDGFR^+^ cells of the human heart(23, 25). These findings demonstrate that the two most discussed putative CPCs do not significantly contribute to cardiomyocytes. When coupled with data demonstrating that most cardiomyocyte renewal can be accounted for by division of pre-existing cardiomyocytes, these findings reduce the likelihood that there is a CPC population in mammalian species.

## ACKNOWLEDGMENTS

We thank Dr. Kyohei Oyamaya for expert training in cardiomyocyte isolation. We thank Dr. Dale Hailey in the Mike and Lynn Garvey Imaging Core within the Institute for Stem Cell and Regenerative Medicine for assistance with fluorescent imaging. This research also was supported by the Cell Analysis Facility Flow Cytometry and Imaging Core in the Department of Immunology at the University of Washington.

